# Mapping high resolution, multidimensional phase diagrams of physiological protein condensates

**DOI:** 10.1101/2025.11.24.690258

**Authors:** Tanushree Agarwal, Tomas Sneideris, Fabian Svara, Klavs Jermakovs, Helena Coyle, Seema Qamar, Emanuel Kava, Rob Scrutton, Nicole Pleschka, Priyanka Peres, Gea Cereghetti, Ewa Andrzejewska, Alejandro Diaz-Barreiro, Gaby Palmer, Antonio J. Costa-Filho, Georg Krainer, Tuomas PJ Knowles, Jonathon Nixon-Abell

**Author notes:** Corresponding authors: Jonathon Nixon-Abell, Tuomas PJ Knowles, Georg Krainer.

## Abstract

Biomolecular condensates are membraneless compartments, crucial for organising and regulating diverse cellular processes. Current approaches to study condensate biology either use simplified recombinant protein systems with limited physiological relevance, or complex live-cell models with restricted experimental control and scalability. Here, we present ExVivo PhaseScan, a droplet microfluidics platform that couples mammalian lysate-based reconstitution with scalable analysis to generate high-resolution phase diagrams of physiological protein condensates. We apply this approach to study two complex multicomponent condensate systems, stress granules and nucleoli, and dissect the physicochemical interactions that influence their stability. We further developed a machine learning pipeline to analyse condensate morphology which we use to reveal how mutations in the ALS-linked protein FUS remodels condensate phase landscapes. We identify liquid-to-solid transitions of mutant FUS within stress granules and nucleoli, and show that these transitions can be reversed by RNA aptamer-based interventions. Together, these findings establish ExVivo PhaseScan as a versatile tool for dissecting the physicochemical and pathological regulation of condensates, with potential to inform therapeutic strategies for diseases driven by aberrant phase transitions.

## Introduction

Biomolecular condensates are membraneless compartments comprising proteins and nucleic acids organised into dynamic assemblies that facilitate the spatial and temporal control of diverse cellular processes [1, 2]. Condensates form through phase separation, a process that drives dispersed biomolecules to demix into dense and dilute phases, enabling condensates to rapidly remodel in response to changing cellular conditions [3]. When the balance of these processes is disrupted, condensates can undergo aberrant phase transitions from dynamic, liquid-like assemblies into gel-like or aggregated states [4]. Such transitions are increasingly recognised as hallmarks of diseases including amyotrophic lateral sclerosis, frontotemporal dementia, and cancer [5, 6]. Understanding how condensates are formed, maintained, and destabilised is therefore critical to both fundamental and disease-focussed research.

A key challenge in understanding these processes lies in identifying the physical, chemical and biological parameters that govern condensate behaviour. Protein interactions, RNA abundance, ionic strength, and osmolarity all contribute to condensate stability [7–10]. Mapping how these variables drive assembly and disassembly typically involves the generation of phase diagrams that capture the conditions under which condensates transition between phase states [11]. While such diagrams are straightforward to obtain in minimal recombinant systems, comprising one or two components, these assays oversimplify the complexity of native condensates, omitting the contributions of cofactors, nucleic acids, and the competing or cooperative interactions among client proteins. Conversely, studies in intact cells capture physiological complexity but lack the experimental control, scalability, and interpretability needed for systematic analysis. Consequently, there remains a need for approaches that combine the biochemical richness of cellular environments with the quantitative precision of *in vitro* systems.

To address this gap, we developed ExVivo PhaseScan, a droplet-based microfluidic platform that uses mammalian cell lysate reconstitution to map condensate phase behaviour at high resolution. This approach combines the compositional complexity of cellular environments with the precision and throughput of microfluidics, enabling robust analysis of multicomponent condensates under near-physiological conditions. We apply ExVivo PhaseScan to stress granules and nucleoli, model condensates involved in stress responses and ribosome biogenesis [13, 14]. Using this platform, we generate high-resolution phase diagrams of these native condensates, dissect the physicochemical interactions driving their assembly, and reveal co-condensation of client proteins that are poorly captured in simplified recombinant systems. Moreover, we demonstrate that disease-associated FUS mutations [15] alter the phase boundaries of both condensate systems, promoting pathological solidification, which we show can be reversed by targeted RNA aptamers. Together, these results demonstrate that ExVivo PhaseScan enables systematic investigation of the physical principles and disease-related perturbations of biomolecular condensates and highlight its potential as a powerful screening platform for identifying therapeutic strategies that restore normal condensate dynamics.

## Results

### Rationale of ExVivo PhaseScan

ExVivo PhaseScan builds on the PhaseScan technology [3], a droplet-based microfluidic approach that generates high-resolution phase diagrams from thousands of microdroplets, enabling rapid exploration of condensate behaviour across large parameter spaces. In ExVivo PhaseScan, we adapt this approach to lysate-based reconstitution of physiological condensates. Here, condensates are reconstituted within microdroplets directly from mammalian cell lysates through the addition of recombinant “scaffold” proteins that nucleate condensate formation and recruit endogenous components from the lysate. The ExVivo PhaseScan platform thus enables quantitative, high-resolution mapping of native-like condensates under near-physiological conditions (Figure 1a).

**Figure 1:**
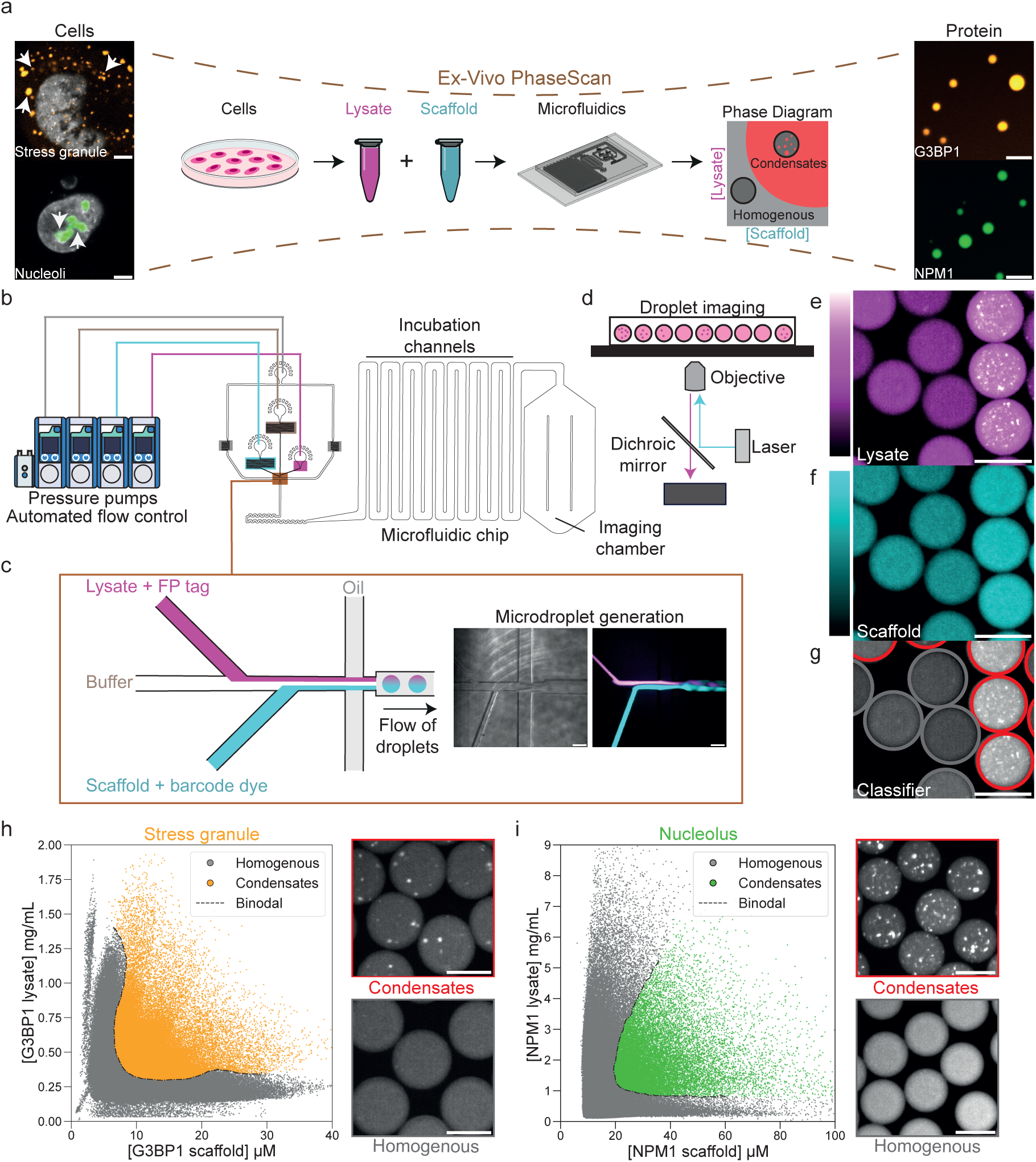
ExVivo PhaseScan generates high resolution phase diagrams of physiological protein condensates. **(a)** Ex-Vivo PhaseScan (centre) exploits the relative strengths of both *in vitro* recombinant protein systems and *in vivo* cellular systems. Left: confocal images from cells expressing fluorescent markers of stress granules (G3BP1-mScarlet) and nucleoli (NPM1-mScarlet). Nuclei are labelled with SPY650 (grey). White arrows indicate condensates. Right: 10 μM AF647 labelled G3BP1 and NPM1 protein condensates formed in 5% w/v PEG (20 kDa). Scale bars, 4 μm. **(b–g)** Schematic of PhaseScan: the microfluidic device is regulated by automated pressure pumps (b). to form microdroplets with varying scaffold and lysate concentrations. The labelled aqueous solutions mix under laminar flow at the microdroplet generating junction, displayed by brightfield (left) and fluorescence (right) microscopy (c). Scale bars, 100 μm. Formed microdroplets are passed down incubation channels before being imaged by widefield microscopy under continuous flow (d). Imaging of both the lysate (e) and scaffold (f) enables quantitative assessment of their relative concentrations. Micrographs here show droplets containing mScarlet-G3BP1 lysate and AF647-labelled G3BP1 scaffold, and subsequent classification of the droplets into ‘condensate’ (red circles) or ‘homogenous’ categories (g). Scale bars, 100 μm. **(h–i)** High resolution phase diagrams for stress granules (h) and nucleoli (i). Each datapoint represents an individual microdroplet classified as either homogeneous (grey) or containing condensates (stress granules-yellow, nucleoli-green). Binodal phase boundaries are plotted (dashed line). Representative widefield images of each classification category are displayed alongside. *n* = 155,584 (stress granule); 159,642 (nucleoli) microdroplets. Scale bars, 100 μm.

### Generation of high-resolution phase diagrams of stress granules and nucleoli

We first applied this approach to investigate reconstituted stress granules and nucleoli. Following an established reconstitution pipeline [4], we generated lysate from HeLa cells stably expressing G3BP1-mScarlet or NPM1-mScarlet, respective markers of stress granules and nucleoli. Adding a small amount of purified G3BP1 (for stress granules) or NPM1 (for nucleoli) scaffold nucleates condensate formation, recruiting endogenous components from the lysate, including the respective mScarlet-tagged markers (Supplementary Figure 1a).

We confirmed specificity of the reconstitution by testing heterologous scaffold-lysate combinations. Robust condensation occurred only when scaffold proteins were matched with their corresponding lysates (e.g. G3BP1 scaffold with G3BP1 lysate); neither BSA nor unrelated scaffolds induced condensate formation under equivalent conditions (Supplementary Figure 1b,c). Thus, condensate assembly in this system requires both the identity of the scaffold and the molecular context of the lysate. To assess the material properties of the reconstituted condensates, we performed fluorescence recovery after photobleaching (FRAP). Both reconstituted stress granules and nucleoli displayed rapid fluorescence recovery (Supplementary Figure 1d,e), consistent with their liquid-like dynamics as reported in cells [5, 6].

To generate phase diagrams of our reconstituted condensates, we loaded the lysate and scaffold into PhaseScan (Figure 1b; see Materials and Methods, Section 9) [3]. This yielded tens of thousands of microdroplets spanning a wide range of scaffold and lysate concentrations (Figure 1c). Generated droplets were then imaged under flow using a fluorescence microscope (Figure 1d), with lysate concentrations being reported by the mScarlet tag (Figure 1e), and scaffold protein concentrations determined by a ‘barcoding’ dye diluted in the protein buffer (Figure 1f). This approach allows us to determine the exact concentration of lysate and scaffold protein within each droplet by comparing them to calibration measurements using known barcode concentrations (see Materials and Methods, Section 9). We next used a custom classifier algorithm to assign each microdroplet as either homogeneous or containing condensates based on the mScarlet tagged marker (Figure 1g).

From these data, we constructed high-density phase diagrams for both condensate types. Stress granules formed robustly over G3BP1 scaffold concentrations of ∼3–40 µM and lysate concentrations of ∼0.1–1.75 mg/mL (Figure 1h), while nucleoli formed over NPM1 scaffold concentrations of ∼8–100 µM and lysate concentrations of ∼0.1–8 mg/mL (Figure 1i). In both cases, sharp binodal boundaries separated homogeneous and phase-separated regions, providing a quantitative map of the conditions under which condensates assemble. Replicate experiments confirmed that the phase boundaries were highly reproducible, with minimal variation between experiments (Supplementary Figure 1f). Importantly, recombinant scaffolds alone did not undergo condensation under the same conditions (Supplementary Figure 1g,h), underscoring the essential role of lysate-derived components.

These results establish ExVivo PhaseScan as a novel platform for generating reproducible, high-resolution phase diagrams of complex near-physiological condensates.

### Mapping the molecular interactions driving stress granule condensation

Having established ExVivo PhaseScan as a robust system for studying the phase behaviour of physiological condensates, we next sought to interrogate the molecular interactions that govern their stability. A central advantage of ExVivo PhaseScan is that it allows for the precise, systematic modulation of physicochemical parameters within a native molecular context. Such perturbations are straightforward *in vitro* but effectively impossible to achieve in living cells, where these parameters cannot be precisely controlled or measured. ExVivo PhaseScan thus provides a unique opportunity to determine the interactions that stabilise or dissolve native condensates.

Stress granules are believed to be maintained by a combination of multivalent, electrostatic, and hydrophobic interactions [7, 8]. We tested how three well-characterised physicochemical modulators of these interactions (poly(A) RNA, varied KCl concentrations, and 1,6-hexanediol) influence stress granule condensation (Figure 2a).

**Figure 2:**
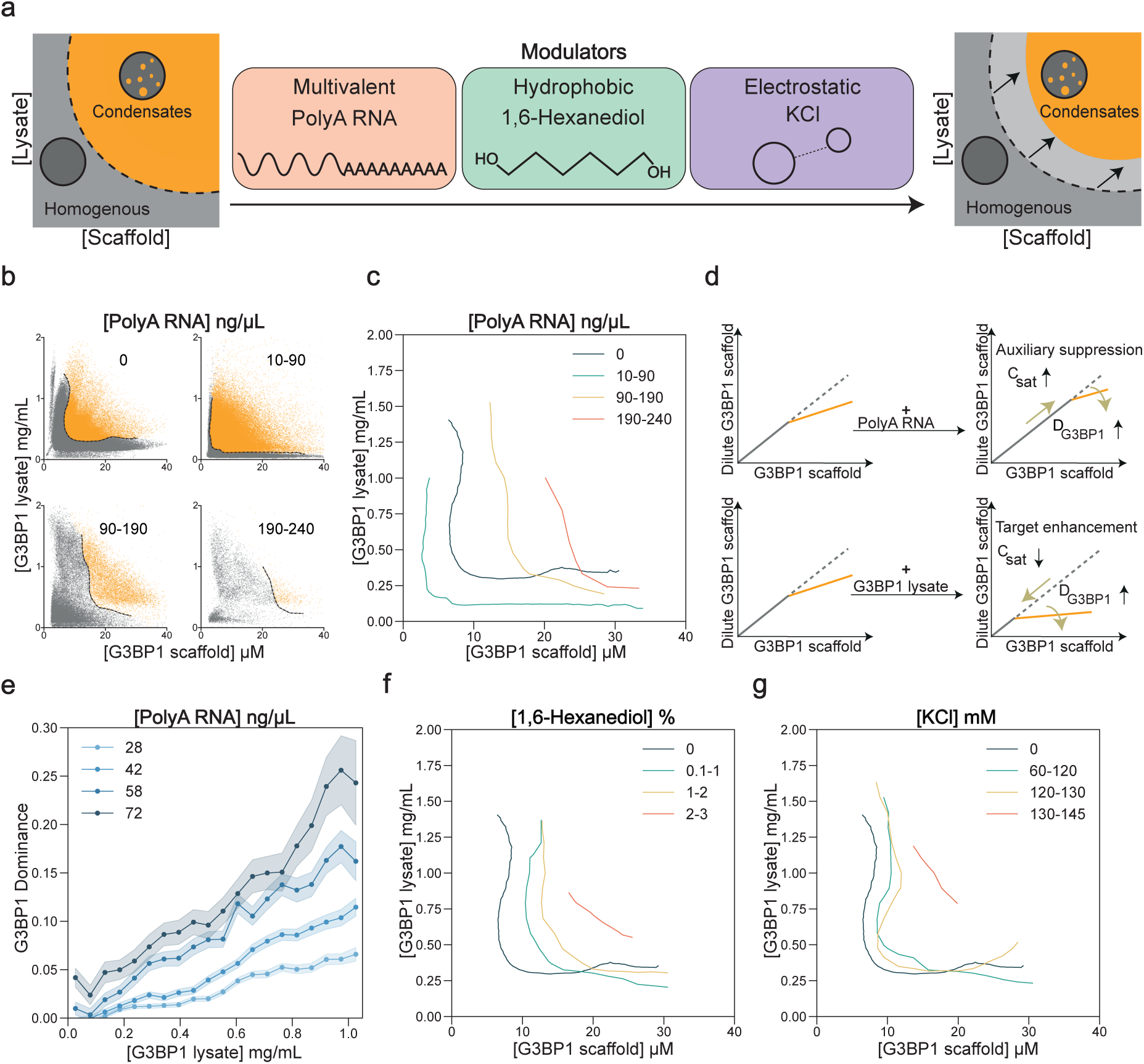
Determining the physiochemical interactions driving stress granule formation. **(a)** Schematic of putative interactions underlying stress granule condensation, and their corresponding physicochemical modulators: Poly(A) RNA, 1,6-Hexanediol, and KCl. **(b)** Phase diagrams showing the effect of increasing Poly(A) RNA concentration on stress granule condensation. Each microdroplet is classed as homogeneous (grey) or containing condensates (yellow), with the binodal phase boundary plotted as a dashed line. *n* =155,584 (0 ng/µL); 298,218 (10–90 ng/µL); 46,256 (90–190 ng/µL); 6,410 (190–240 ng/µL) microdroplets. **(c)** The phase boundaries as shown in (b) plotted together. **(d)** Schematic of two representative mechanisms inferred from dominance analysis, auxiliary suppression (top panel) and target enhancement (bottom panel). C_sat_: saturation concentration. D_G3BP1_: G3BP1 dominance. **(e)** G3BP1 dominance plotted as a function of G3BP1 lysate concentration for four different concentrations of PolyA RNA (28, 42, 58, and 72 ng/μL). **(f–g)** Extracted phase boundaries demonstrating the effect of varying 1,6-Hexanediol (f) concentrations (0.1–3 %) and KCl (g) concentrations (60–145 mM) on stress granule condensation.

### Poly(A) RNA tunes stress granule condensation in a biphasic manner

We first titrated poly(A) RNA across a broad range of concentrations and constructed phase diagrams for each condition (Figure 2b). At low RNA concentrations (10–90 ng/µL), stress granule condensation was enhanced. This is likely driven by the capacity of RNA to promote multivalent interactions between RNA-binding proteins enriched in stress granules [7]. At intermediate concentrations (90–240 ng/µL) condensates progressively dissolved, displaying a re-entrant transition [9], likely due to the disruption of intermolecular protein interactions or increased net charge repulsion, as observed previously in recombinant G3BP1 condensates [10, 11]. At higher RNA concentrations the system became entirely homogeneous (Supplementary Figure 2a). Plotting the binodal phase boundaries at varying RNA concentrations highlights this biphasic behaviour (Figure 2c), suggesting an important dual role for RNA in modulating stress granule condensation.

To mechanistically interpret these changes, we applied a dominance analysis framework [12] that partitions the relative energetic contributions of different components to condensation. Rather than tracking whether condensates form, this method quantifies how scaffold saturation concentration (C_sat_) shifts under different conditions and attributes those shifts to changes in the contribution of specific interactions. In practical terms, this allows us to ask whether condensation is primarily driven by scaffold interactions or by additional lysate-derived factors, and how RNA alters this balance (see Supplementary Text, Section 1.1 and Supplementary Figure 2b). We identified two distinct mechanisms during stress granule condensation. (1) Auxiliary suppression (Figure 2d, top panel), where for RNA concentrations above 28ng/ul, a further increase in RNA concentration raised the scaffold saturation concentration (C_sat_) while increasing scaffold dominance (D_G3BP1_), meaning that non-scaffold interactions contribute less to the overall free energy of condensation (2) Target enhancement (Figure 2d, bottom panel), where increasing lysate concentration lowered C_sat_ while also increasing D_G3BP1_, indicating that scaffold interactions became more important to the free energy of condensation as lysate levels rose. Figure 2e represents the increase in scaffold dominance as a function of both lysate and RNA concentrations. The dominance framework analysis not only supports previous *in vitro* observations that RNA’s dissolution effect is driven by mitigating non-scaffold interactions [13], but also reveals that stress granule formation energetics are governed by promiscuous G3BP1 interactions with multiple lysate components - binding promiscuity that cannot be fully recapitulated in reductionist *in vitro* systems.

### Hydrophobic interactions promote stress granule stability

We next evaluated the role of hydrophobic interactions in stress granule condensation by introducing 1,6-hexanediol, an aliphatic alcohol known to disrupt weak hydrophobics [14]. Increasing 1,6-hexanediol (0.2–6%) caused a progressive retreat of the phase boundary towards the homogeneous region, with complete dissolution above ∼2% (Figure 2f; Supplementary Figure 3a). This suggests that hydrophobic interactions, likely arising from transient side-chain contacts (e.g. π–π, cation–π) and aliphatic packing [15] within intrinsically disordered regions of stress granule proteins, are important cohesive forces in stress granule condensation.

### Charge shielding disrupts electrostatic cohesive forces in stress granules

Finally, we investigated electrostatic contributions by titrating KCl across a physiologically relevant concentration range (60–160 mM). Increasing ionic strength progressively dissolved stress granules, with complete loss of condensates at concentrations above 145 mM (Figure 2g; Supplementary Figure 3b). This effect likely reflects charge shielding of RNA–protein interactions which has been suggested to play an important role in stress granule nucleation and stability in yeast and mammalian cells [16], reinforcing the concept that electrostatic interactions are a central cohesive force in stress granules.

Together, these data reveal how distinct classes of physicochemical interactions contribute to stress granule stability. Poly(A) RNA promotes condensation at low concentrations but disrupts it at high concentrations, while hydrophobic and electrostatic interactions are essential stabilising forces whose disruption leads to stress granule dissolution. Beyond providing mechanistic insights into stress granule stability, these findings illustrate the potential of ExVivo PhaseScan in providing a quantitative framework for mapping how physicochemical parameters shape the phase behaviour of numerous other native physiological condensates.

### ExVivo PhaseScan enables condensate client profiling

A key advantage provided by ExVivo PhaseScan is the ability to study physiologically relevant clients recruited into lysate-reconstituted condensates; interactions often absent in minimal recombinant assays due to the simplistic nature of single component condensate proxies. We chose to explore this in the context of stress granules due to their recruitment of diverse client proteins [17–19] and because prior work has shown that the proteome of lysate-reconstituted stress granules closely mirrors that of their *in cellula* counterparts [4].

We first compared the incorporation of three stress granule clients, Fused in Sarcoma (FUS) [17], interleukin-38 (IL-38) [18], and pyruvate kinase M2 (PKM2) [19, 20] into cellular, lysate-based, and recombinant stress granule systems. To confirm that each client is indeed recruited to cellular stress granules, we expressed tagged versions of each client in a HeLa line stably expressing a stress granule marker (G3BP1-mScarlet). Upon stress induction, G3BP1-positive puncta indicated the formation of stress granules, to which each client was robustly recruited (Figure 3a).

**Figure 3:**
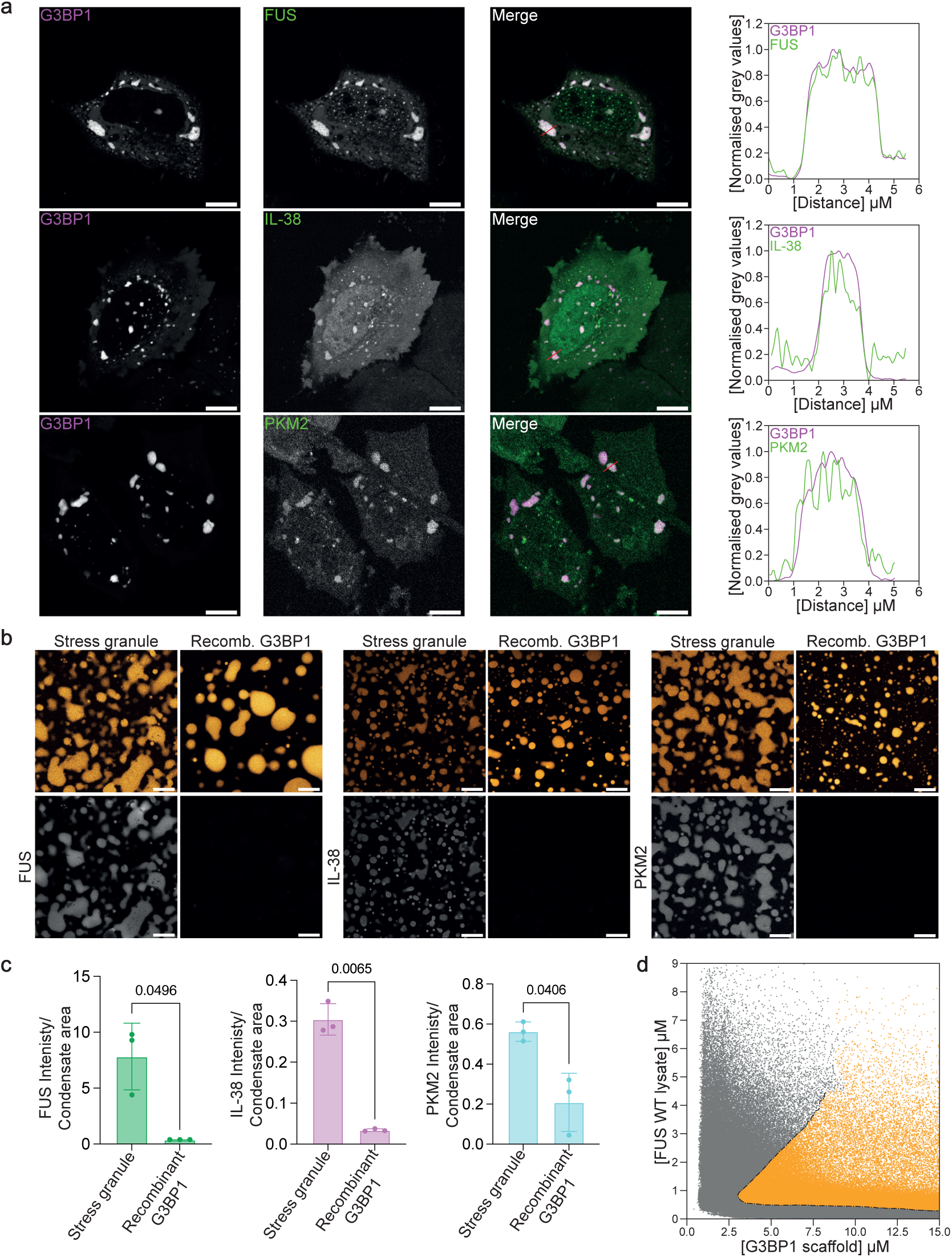
Ex-Vivo PhaseScan enables the study of physiological stress granule-associated clients. **(a)** Confocal images of HeLa cells under stress conditions (see Materials and Methods, Section 5) stably expressing G3BP1-mScarlet and either mEmerald tagged FUS, IL-38, or HA tagged PKM2 which was visualised using an mEGFP-anti-HA-frankenbody. The corresponding line scan plots are graphed from the red lines on the merge panels. Scale bars, 10 μm. **(b)** Confocal images of lysate-reconstituted stress granules (15 µM G3BP1 + 1 mg/mL mScarlet-G3BP1 lysate) and recombinant-only G3BP1 condensates (15 µM AF647-G3BP1 + 10 ng/µL PolyA RNA) in the top row. Bottom row: client recruitment of 1 µM FUS-GFP, 1 µM AF488-IL-38, 1 µM AF488-PKM2. Scale bars, 20 μm. **(c)** Quantification of the client fluorescent intensity per condensate area for lysate-based vs recombinant conditions as shown in (b). Mean ± s.d., *n* = 3 independent experiments. Unpaired two-tailed t-test with Welch’s correction (*p*-values reported on the figure). **(d)** PhaseScan of FUS-mEmerald lysate seeded with G3BP1 scaffold; microdroplets classified as homogeneous (grey) or condensate-containing (yellow), with the binodal phase boundary plotted as a dashed line. *n* = 397,104 microdroplets. Panel duplicated as Figure 4f for disease-focused analyses.

We next tested whether client recruitment was recapitulated in our lysate-based reconstitution system and how this compared to recombinant G3BP1 condensates. We reconstituted stress granules using 15 µM G3BP1 and 1 mg/mL lysate generated from G3BP1-mScarlet cells. Recombinant-only condensates were generated by combining 15 µM G3BP1 with 10 ng/µL poly(A) RNA. Both approaches robustly produced condensates (Figure 3b, top row). However, upon addition of fluorescently labelled client proteins (Figure 3b, bottom row), only lysate-based condensates showed client enrichment, whereas recombinant condensates failed to recapitulate the client recruitment observed in cells (Figure 3c).

We next demonstrated that this client recruitment is retained in ExVivo PhaseScan by complementing FUS-mEmerald lysate with G3BP1 scaffold protein (Figure 3d). FUS efficiently partitioned into G3BP1 condensates confirming that the approach can simultaneously resolve phase boundaries and client recruitment with high spatial and quantitative precision. Together, these data demonstrate that ExVivo PhaseScan enables quantitative mapping of condensate phase behaviour while directly capturing client recruitment within a native-like biochemical environment.

### ALS-linked FUS mutations remodel condensate phase landscapes and are reversible by RNA aptamers

We next asked how amyotrophic lateral sclerosis (ALS)-linked FUS variants reshape these recruitment landscapes and condensate biophysical states. Pathogenic mutations in FUS drive aberrant phase transitions linked to ALS [21, 22], yet understanding these transitions in a physiologically relevant setting has been challenging: recombinant assays miss native cofactors, whereas cellular experiments lack the control and throughput needed to chart reliable phase boundaries. Under basal conditions wildtype FUS displays a broad nuclear localisation but translocates to stress granules or nucleoli under stress conditions or when harbouring certain ALS mutations [23]. Here we use ExVivo PhaseScan to quantify how ALS-linked FUS variants reshape condensate phase landscapes in nucleoli and stress-granules, and whether these pathological states can be reversed.

We focused on two ALS-linked FUS variants that perturb condensate behaviour through distinct mechanisms and in different subcellular contexts. The G156E mutation lies within the prion-like low-complexity domain and increases self-association, favouring gelled or aggregated states in the nucleus where FUS engages nucleolar components and influences nucleolar architecture [24] [25]. In contrast, R522G disrupts the PY-NLS that mediates transportin/Kapβ2-dependent nuclear import, causing cytoplasmic mislocalisation and enhanced association with stress granules [26]. These mutants therefore represent orthogonal disease mechanisms that manifest as alterations in compartment-specific phase transitions (Figure 4a).

**Figure 4:**
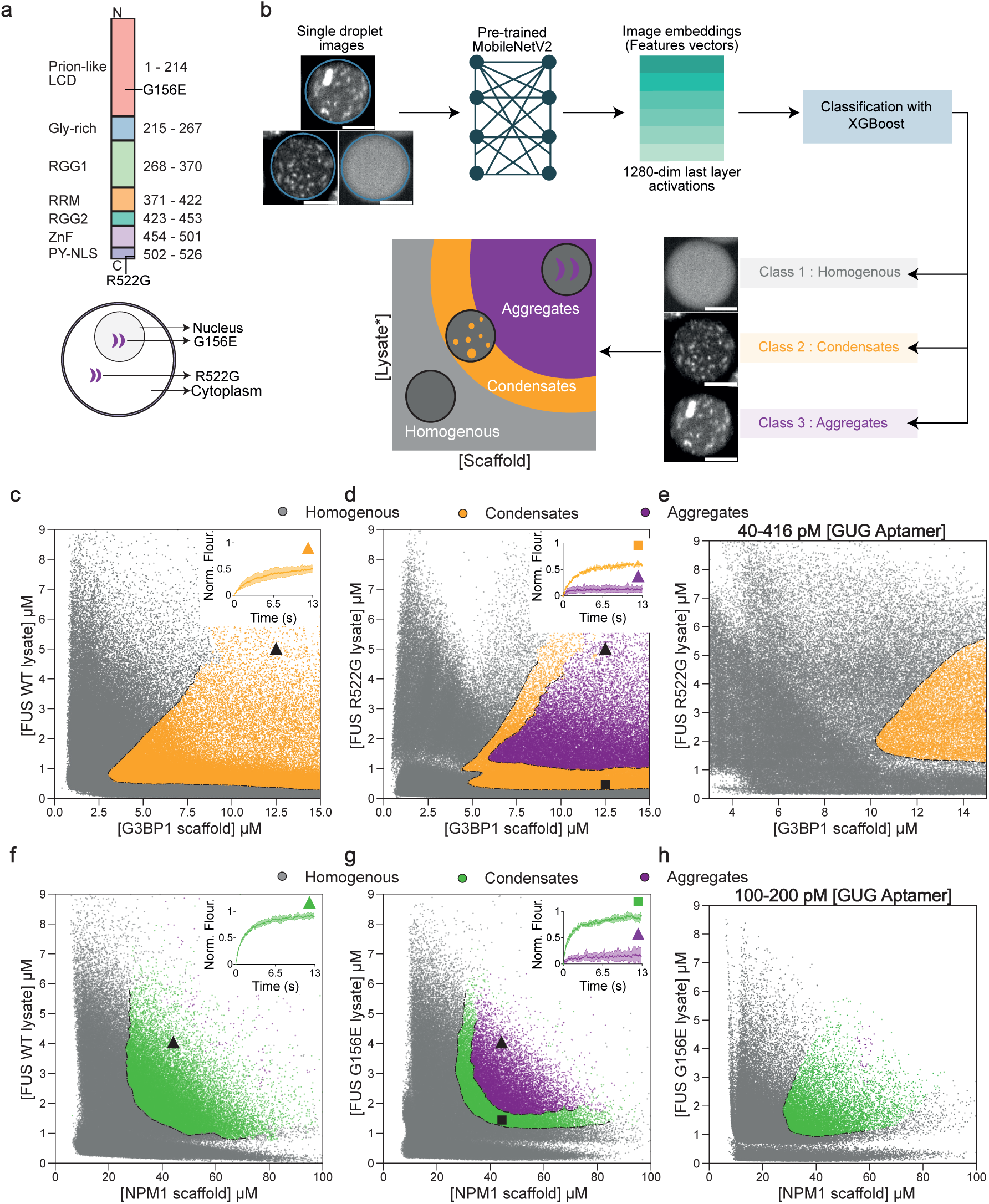
ALS-linked FUS mutations alter condensate phase behaviour and are rescued by RNA aptamers. **(a)** Top: Schematic of FUS domain organisation with key mutations highlighted: G156E in the low-complexity domain (LCD) and R522G in the proline-tyrosine nuclear localisation signal (PY-NLS). Bottom: Compartment-specific effects of these mutations, where G156E promotes FUS nuclear aggregation and R522G drives cytoplasmic mislocalisation. **(b)** Machine learning pipeline for classification of droplet morphologies in Ex-Vivo PhaseScan where representative single-droplet images were processed through a pre-trained MobileNetV2 network to generate 1280-dimensional feature embeddings, which were classified using XGBoost into three morphological classes: homogeneous, condensates, and aggregates for subsequent plotting onto phase diagrams. Scale bars, 50 μm. **(c–e)** PhaseScan of G3BP1 scaffold with FUS-mEmerald WT (c) and R522G (d) lysates. Panel (e) shows how a GUG RNA aptamer alters the R522G phase boundaries. Microdroplets are assigned to homogeneous (grey), condensate (orange) and aggregate (purple) categories, with the binodal phase boundary shown as a dashed line. *n* = 397,104 (WT); 409,859 (R522G); 131,317 (R522G+GUG) microdroplets. Insets show FRAP traces from condensates and aggregates and their corresponding positions on the phase diagram. Mean ± s.d., *n* = 3 independent experiments. **(f–h)** PhaseScan of NPM1 scaffold with FUS-mEmerald WT (f) and R522G (g) lysates. Panel (h) shows how a GUG RNA aptamer alters the R522G phase boundaries. Microdroplets are assigned to homogeneous (grey), condensate (orange) and aggregate (purple) categories, with the phase boundaries shown as a dashed line. *n* = 173,630 (WT); 140,300 (G156E); 93,138 (G156E+GUG) microdroplets. Insets show FRAP traces from condensates and aggregates and their corresponding positions on the phase diagram. Mean ± s.d., *n* = 4 independent experiments. Panel f duplicated from Figure 3d above.

To model these differences in ExVivo PhaseScan, we generated lysates from HeLa cells stably expressing FUS-mEmerald wildtype (WT), G156E or R522G at similar expression levels (Supplementary Figure 4a). We paired each lysate with its corresponding scaffold protein, reconstituting nucleoli using NPM1 with the G156E lysate and stress granules using G3BP1 with the R522G lysate. Confocal imaging revealed notable differences in the morphology of WT and mutant FUS condensates. While WT FUS adopted spherical liquid-like condensates, G156E and R522G lysates produced aggregate-like structures with their respective scaffolds (Supplementary Figure 4b). To enable a scalable and unbiased analysis of these changes in ExVivo PhaseScan, we developed an automated machine learning pipeline to classify each microdroplet image as homogeneous, condensate-, or aggregate-containing (Figure 4b; see Supplementary Text, Section 2.1 and Materials and Methods, Section 10). To quantitatively assess classifier performance, confusion matrices were generated for three independently trained models (Supplementary Figures 4c-e). All models exhibited high precision and recall across the three morphological classes (weighted F1 ≈ 97-98%). We also validated that our classifier relies on morphology-based condensate features rather than class-specific information leakage (Supplementary Figure 5a; see Supplementary Text, Section 2.2). This performance highlights the robustness of the pipeline for distinguishing subtle morphological differences, demonstrating its general utility for high-throughput droplet phenotyping.

We subsequently applied the trained model to map how scaffold and lysate concentrations shape phase boundaries for WT and ALS-linked FUS in stress granule and nucleolar reconstitutions. In stress granules WT FUS yielded only homogeneous and condensate droplets across the concentration ranges tested (Figure 4c), with condensates showing typical rapid fluorescence recovery by FRAP (Figure 4c inset panels). By contrast, R522G produced aggregate-assigned droplets emerging at elevated scaffold and lysate concentrations (Figure 4d). Assemblies sampled from this region were immobile by FRAP, consistent with aggregate-like solidification (Figure 4d inset panels).

Building on (i) previous reports [10, 27], (ii) our earlier observation that RNA levels tune stress-granule phase behaviour, and (iii) FUS’s extensive RNA-binding interfaces, we hypothesised that a defined RNA-based intervention could bias assemblies toward reversible, fluid interactions. Consistent with this hypothesis, a FUS-binding GUG RNA aptamer [28] collapsed the aggregated region for R522G and restored a condensate-only landscape (Figure 4e), whereas a scrambled UGU control was inert (Supplementary Figure 5b).

We observed analogous behaviour in the nucleolus. WT FUS again produced only homogeneous and condensate droplets, (Figure 4f), with condensates exhibiting rapid FRAP recovery (Figure 4f inset panels). In contrast, G156E displayed an aggregated region at higher NPM1 and lysate concentrations (Figure 4g); FRAP distinguished these immobile aggregates from liquid-like condensates at lower concentrations (Figure 4g inset panels). The same FUS-binding aptamer eliminated the aggregated region for G156E (Figure 4h), and the scrambled control again had no effect (Supplementary Figure 5c). These data indicate that aptamer-mediated rescue generalises across compartments and mutant mechanisms. Crucially, we generated a G156E recombinant-only condensate which does not display any aggregates (Supplementary Figure 5d), reinforcing the importance of using lysate-based reconstitution over a purely recombinant protein system.

Together, these data show that WT FUS in both stress granules and nucleoli remains liquid-like across broad concentration ranges, whereas ALS-linked G156E and R522G aggregate above critical scaffold/lysate levels. Crucially, RNA aptamer-based intervention can prevent this aggregation, providing a putative strategy for restoring healthy condensate behaviour. ExVivo PhaseScan thus provides a scalable platform to map, compare, and therapeutically probe disease-relevant phase landscapes across proteins, mutations, and modulators.

## Discussion

Our study introduces ExVivo PhaseScan, a lysate-based microfluidic platform that enables quantitative, high-resolution mapping of condensate phase behaviour under near-physiological conditions. Traditional *in vitro* assays using recombinant proteins provide experimental control but lack molecular complexity that strongly influences condensate stability and composition in cells [1, 2, 29]. Conversely, cellular studies preserve physiological context but limit mechanistic interrogation [30]. ExVivo PhaseScan unites these regimes, enabling the precise modulation of biochemical variables while retaining native cofactors, thereby providing a robust framework for dissecting the physical, chemical and biological principles governing both adaptive and pathological phase transitions (Figure 1).

### Quantitative dissection of stress granule assembly and client profiling

Using stress granules as a test case, we show through systematic perturbation with poly(A) RNA, KCl, and 1,6-hexanediol that their condensation depends on a finely tuned balance of electrostatic, hydrophobic, and multivalent interactions (Figure 2). RNA stoichiometry emerged as a principal regulator of stress granule phase behaviour. Low RNA concentrations enhanced condensation, whereas high RNA levels solubilised assemblies. This biphasic response is supported by previous *in vitro* studies showing that RNA can act as both a multivalent cross-linker and a solubilising agent depending on valency and concentration [10, 11, 31, 32] These data raise the intriguing possibility that RNA stoichiometry may act as a molecular rheostat [33, 34] balancing stress granule formation and dissolution during stress and recovery. In further support of this model, our dominance analysis revealed that the RNA re-entrant transition is not in fact due to mitigating G3BP1’s energetic contribution to stress granule formation, but instead is mediated through a reduction in the energetic contributions from non-scaffold interactions. In addition to RNA, we show that perturbations with 1,6-hexanediol and KCl indicate that weak hydrophobic interactions provide structural cohesion, while electrostatic interactions are a key cohesive force governing stress granule stability. These data align with previous work that demonstrates the relative significance of hydrophobic and electrostatic interactions in recombinant G3BP1 condensates [13, 14, 27]. Together, we have, for the first time, dissected the forces stabilising stress granule assembly in near-physiological condensates, providing insight into how biologically relevant fluctuations in these parameters can fine-tune condensate stability and dynamics.

Another key advantage of our approach is its capacity to uncover client recruitment logic that is largely lost in minimal systems. While recombinant G3BP1 condensates are able to recruit some native clients [27], we show that they fail to capture many others including FUS, IL-38, and PKM2. By contrast lysate-based condensates robustly concentrated these clients, indicating that their partitioning depends on context-dependent interactions beyond the scaffold itself (Figure 3). These likely include complementary charge patterns, low-affinity hydrophobic contacts, and protein–RNA-mediated bridging, which are absent in minimal systems [35].Importantly, these are the same interactions that, as discussed above, are relevant for stress granule stability, suggesting that condensate composition and phase boundaries are co-regulated by the same interaction parameters. Together, these results illustrate the unique capacity of ExVivo PhaseScan to link phase boundaries with selective molecular partitioning in the same experimental system, and to explore how physicochemical modulators influence complex multicomponent physiological condensates.

### Pathological insights into stress granule and nucleolus phase transitions

Beyond mechanistic characterisation, ExVivo PhaseScan provides a powerful platform for exploring the molecular origins of pathological phase transitions. Here we quantify how ALS-linked FUS mutants perturb phase equilibria in their native biochemical environments (Figure 4). The R522G and G156E variants are respectively associated with impaired nucleocytoplasmic FUS transport and aberrant condensation inside cells [24, 26]. Mapping high resolution phase diagrams, we identified distinct, compartment-specific liquid-to-solid transitions: R522G drove aggregation within stress granule reconstitutions, while G156E induced aggregation within nucleolar reconstitutions. To enable this analysis, we developed a machine learning–based morphological classifier to enable the unbiased, high-throughput quantification of droplet states, distinguishing liquid-like condensates from solid aggregates across tens of thousands of reaction compartments. This quantitative mapping directly links pathogenic mutations to altered phase equilibria and material states of physiological condensates. To our knowledge, this is the first example of high resolution phase diagrams detailing both liquid and aggregate phase transitions.

Strikingly, we show that a FUS-binding RNA aptamer can reverse the pathological aggregate transition, restoring liquid-like condensate behaviour, while a scrambled control was inert. Structured RNAs have previously been shown to buffer aberrant FUS condensation and toxicity [36] supporting a therapeutic model wherein RNA binders can be used to modulate condensate material properties. Our results extend this concept to illustrate the potential of ExVivo PhaseScan for high-throughput screens of RNA and pharmacological phase modulators in native biochemical environments, establishing the approach as a powerful tool for identifying molecules that restore liquid-like behaviour in disease-associated assemblies.

### Future directions & conclusions

The methodological flexibility and quantitative power of ExVivo PhaseScan open several exciting avenues for future research. We envisage extending this approach to additional condensate types, including P-bodies, nuclear speckles, paraspeckles, germ granules, and disease-associated assemblies to uncover both shared molecular principles and condensate-specific rules. In this context, higher-dimensional phase mapping incorporating additional biochemical variables such as post-translational modifications, cofactors, or chaperone activity could further resolve how regulatory factors shape condensate energetics and composition. A promising next step would be to couple ExVivo PhaseScan with droplet-sorted proteomic or transcriptomic analyses, allowing molecular deconvolution of phase boundaries and identification of specific proteins, RNAs, or modifications that shift phase equilibria or modulate condensate composition. Finally, as exemplified above, the automated and scalable nature of the assay makes it well-suited for screening applications, enabling quantitative evaluation of small molecules, peptides, or nucleic acids that restore or modulate condensate dynamics in native systems. To our knowledge, there are no currently existing approaches that facilitate this type of screen on complex multicomponent condensates.

In summary, ExVivo PhaseScan provides a system for mapping the phase behaviour of multicomponent condensates under physiologically relevant conditions. By combining lysate-based reconstitution with microfluidics and automated image analysis, it enables the simultaneous characterisation of phase boundaries, client recruitment, and material states across broad parameter spaces. Our finding that disease-associated liquid-to-solid transitions can be reversed by RNA aptamers underscores its potential to guide targeted therapeutic strategies. More broadly, this approach offers a scalable and generalisable platform for elucidating the physical, chemical and biological principles of biomolecular condensation and for probing how their dysregulation drives pathology.

## Materials and Methods

### 1. Plasmids and cloning

Constructs for protein expression encoding G3BP1 (UniprotID:Q13283) and NPM1 (UniprotID: P06748) were sub-cloned by PCR amplification from HeLa cell cDNA for G3BP1 (SQ_001-002) and a GFP-NPM1 plasmid from the Wang lab (Addgene plasmid #17578) for NPM1 (SQ_003-004). Each amplicon was cloned into the bacterial expression vector pOPINS (Merck) containing a ULP protease cleavable N-terminal His-SUMO Tag which was linearised using primers SQ_005-006. For constructs used to generate stale lines, transgenes were cloned from various source plasmids into a custom PiggyBac vector (P455) digested with *BsrGI-HF* and *Mlu1-HF* (NEB). The resultant PiggyBac (PB) plasmids included PB_mScarlet-G3BP1 (P376; JNA_405-408), PB_mScarlet-NPM1 (P378; JNA_405,409-411) and PB_mEmerald-FUS WT (P379; JNA_396-399), G156E (P382; JNA_396-399), R522G (P381; JNA_399,402-404) variants. For transient expression constructs, FUS-mEmerald (P560) was generated by PCR amplification of FUS from an existing mCherry-FUS construct (P281; JNA_905,906) and cloned into an N1-mEmerald expression vector. Primers JNA_035 and JNA_907 were used to amplify the N1-mEmerald backbone. IL-38-mEmerald (P562) was made using a synthesised gene block (IDT) encoding IL-38 into the N1-mEmerald backbone amplified as above. PKM2-HA (P566) was generated by PCR amplification of PKM2 from an existing source plasmid (P561) with an HA tag incorporated in the reverse primer (JNA_929, 930). The HA-frankenbody plasmid (Addgene plasmid #129590) was a kind gift from the Tim Stasevich Lab. All cloning procedures were carried out by PCR amplification using Q5 polymerase (NEB) followed by Gibson Assembly (NEB). Plasmid sequences were verified by Oxford Nanopore sequencing (Plasmidsaurus). The complete list of primers is provided in Table 1.

**Table 1:**
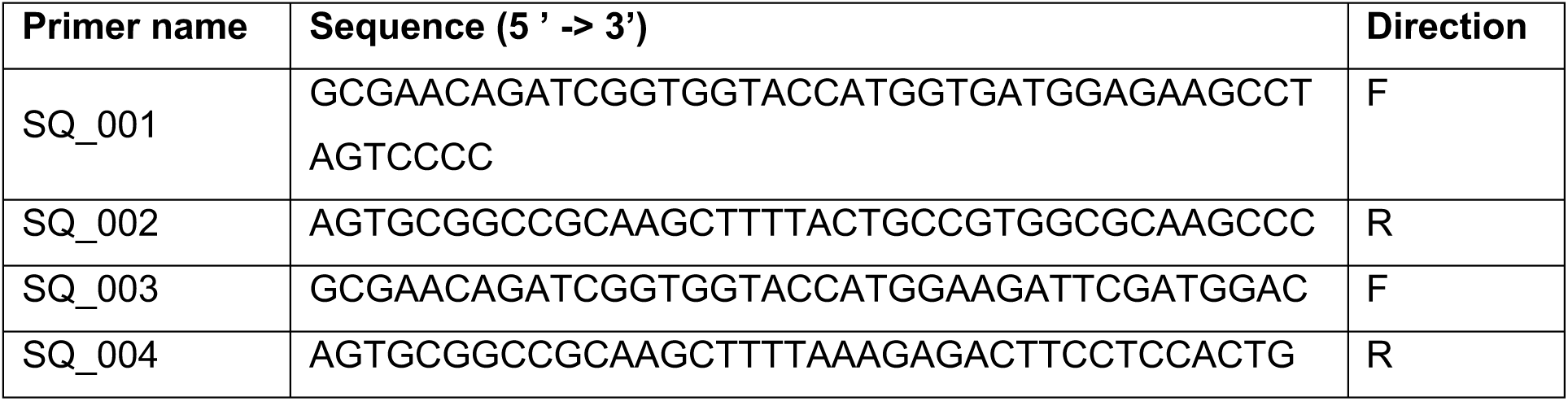
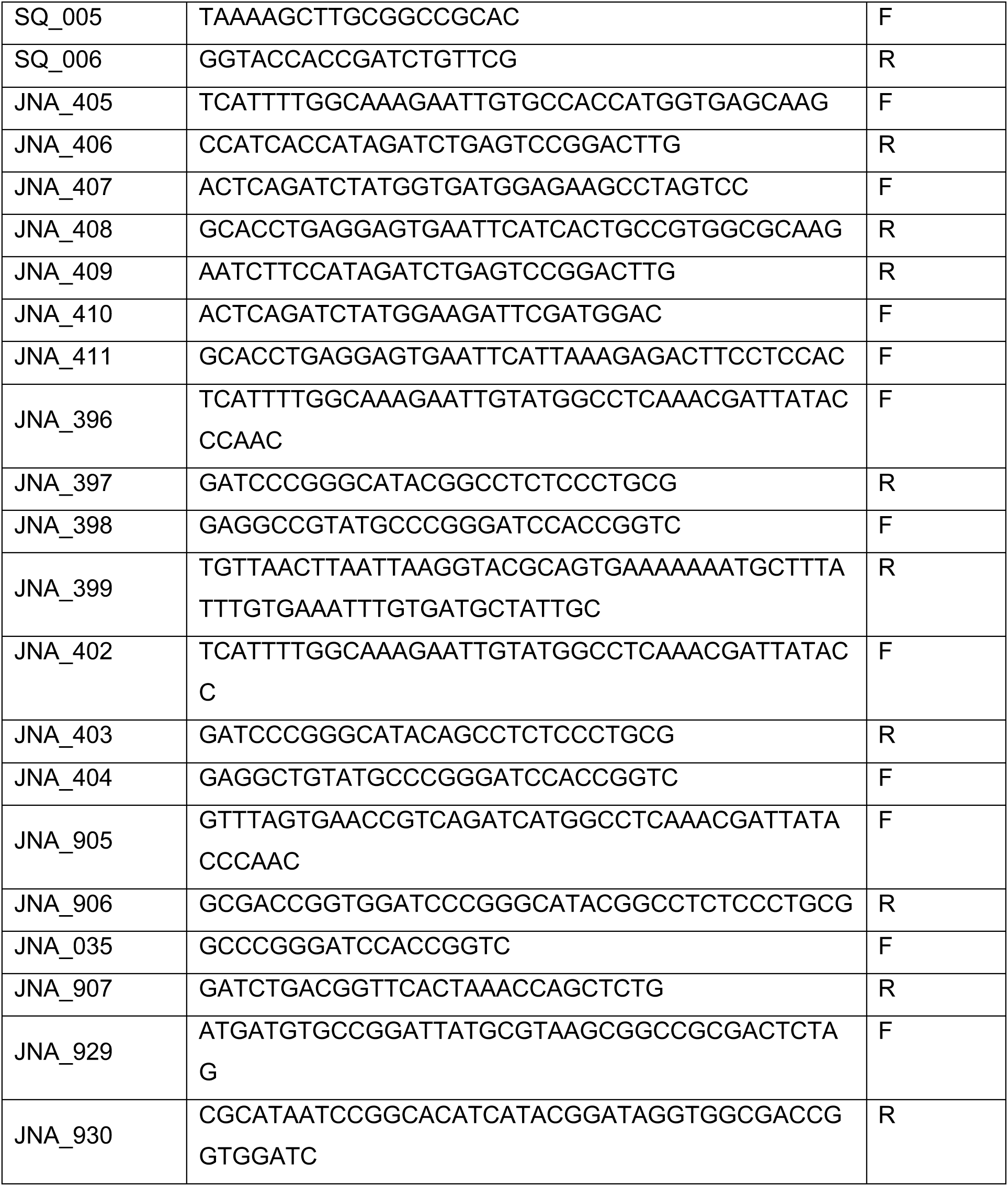
Primers used for cloning.

### 2. Protein purification and labelling

His-SUMO tagged pOPINS constructs for G3BP1 and NPM1 expression were transformed into competent *E. coli* BL21 (DE3) cells (NEB) and expressed overnight in TB autoinduction media (Formedium). Cells were harvested by centrifugation and lysed using a high-pressure cell disruption system (Constant Systems). The resulting lysate was clarified by ultracentrifugation at 100,000 × *g* to remove cell debris and subsequently loaded onto an Ni-Sepharose Advance column (Bioserve) for protein isolation. Protein purification was carried out using a standard Ni-affinity protocol. Eluates containing the target protein were pooled, incubated with ULP protease to remove the His-SUMO tags, and dialyzed against dilution buffer (50 mM HEPES, 200 mM NaCl, 1 mM DTT, pH 7.5). Following tag cleavage, the sample was passed over a second Ni-Sepharose Advance column to remove the His-SUMO tag and subsequently purified by size-exclusion chromatography on a Superdex 200 Increase column (Cytiva) equilibrated with storage buffer containing 50 mM HEPES, 400 mM NaCl, 1 mM DTT, pH 7.5. Purified protein fractions were pooled, concentrated, aliquoted, snap-frozen in liquid nitrogen, and stored at –80 °C until use. For IL-38, *E. coli* BL21 (DE3) cells (NEB) were transformed with a plasmid expressing wild-type human IL-38 (Uniprot: Q8WWZ1) and grown in TB media until OD_600_ = 0.6. Then, IPTG was added to a final concentration of 500 mM and cells were grown at 22 °C overnight. After pelleting by centrifugation, cells were resuspended in lysis buffer (50 mM Tris pH 8, 500 mM NaCl, 10 mM β-ME, 0.5% Triton X-100) supplemented with protease inhibitor tablets and lysed by sonication. Lysates were cleared by centrifugation and supernatant was used for purification using Ni-NTA resin following manufacturer’s instructions. Washes were performed with lysis buffer supplemented with 20 mM imidazole and eluted in 50 mM Tris pH 8.0, 500 mM NaCl, 2 mM β-ME, 5 % glycerol, and 200 mM imidazole. Fractions were dialyzed overnight into 50 mM HEPES pH 7.0, 50 mM NaCl, 5 % glycerol and 2 mM β-ME. Purity was confirmed by SDS-PAGE and Coomassie blue staining. Pure protein was concentrated using a 10 kDa Amicon concentrator to 1 mg/ml, aliquoted, and frozen at -80°C. For PKM2, *E. coli* Rosetta (DE3) cells (Sigma) were transformed with a plasmid expressing wild-type human PKM2 (Uniprot: P14618). Cells were grown at 37 °C in LB media containing 30 mg/ml chloramphenicol and 100 mg/ml carbenicillin until reaching OD_600_ = 0.6. Then, IPTG was added to a final concentration of 0.1 mM. Cells were grown at 16 °C for 12 h, harvested by centrifugation, and resuspended in ice cold purification buffer (100 mM Tris-HCl pH 7.4, 200 mM NaCl, 1 mM MgCl_2_, 10 % glycerol, 1 mM PMSF, 1 mM DTT) supplemented with protease inhibitor tablets and lysed by sonication. After clearing the lysates by centrifugation, the supernatant was loaded on a pre-packed Strep-Tactin Superflow Plus column following the manufacturer’s instructions. Proteins were eluted using purification buffer supplemented with 2.5 mM D-desthiobiotin. Purity was confirmed by SDS-PAGE and Coomassie blue staining, and pure aliquots were stored at -80°C until further use. FUS-GFP WT and G156E were a kind gift from the St George-Hyslop lab and were purified as described previously [3, 37]. Proteins were labelled with AlexaFluor 647 and AlexaFluor 488 NHS ester (Thermo Fisher Scientific) according to the manufacturers instructions. Proteins were incubated with dye at 4°C overnight, and unreacted dye was subsequently removed using appropriate spin columns (Sigma Aldrich).

### 3. Cell line generation and maintenance

Stable HeLa cell lines were generated expressing target genes appended with fluorescently tagged proteins using a Piggybac based integration system. Genes of interest were cloned as described above into a custom PiggyBac vector. Constructs were transfected into HeLa cells alongside a transposase plasmid to induce integration using a 3µl: 1 µg: 0.5 µg ratio of Fugene : PB plasmid : transposase plasmid. Fluorescence-activated cell sorting (FACS) was used to select positively integrated cells 3 days post transfection, with a second sort 6 days later confirming 99% purity. HeLa lines were cultured in DMEM (Gibco) supplemented with 10% FBS (Gibco) and 1% GlutaMax (Thermo) and maintained at 37 °C in 5% CO_2_.

### 4. Lysate Generation

Lysates were generated from the G3BP1-mScarlet, NPM1-mScarlet, FUS-mEm WT, FUS-mEm G156E and FUS-mEm R522G cell lines mentioned above. Cells were grown in 10 cm cell culture dishes to 100% confluency. At confluency, cells were briefly washed in 10ml ice cold PBS (Gibco) prior to being scraped and harvested on ice in 5ml fresh PBS. Cells were pelleted at 300 g for 3-5 minutes. And stored at –80°C until further use. 250 µL of ice-cold lysis buffer (50mM Tris buffer pH 7.4, 0.5% NP40 (Sigma), 1 mini protease inhibitor tablet (Roche)) was mixed with 6.25 µL of RNAse inhibitor (NEB) and added to each pellet. Lysates were mixed and allowed to stand at room temperature for 3 minutes, before being cleared (5 minutes at 21, 000 g). Supernatants were transferred to fresh Eppendorf tubes and the concentration of each lysate was measured using a DCA protein assay kit (Bio-Rad). Lysates of FUS-mEmerald WT, G156E and R522G (8 mg/ml) were dispensed in triplicate into black, flat-bottom 384-well plates (Corning) and their comparative fluorescence intensities were measured using a CLARIOstar plate reader (BMG Labtech).

### 5. Live cell imaging

For imaging experiments cells were seeded in triplicate into Matrigel-coated 8-well glass bottom chambered coverslips (Ibidi, 80826). At ∼70% cell confluency, cells were transiently transfected using FuGene HD (Promega) and 150 ng of the indicated plasmid according to the manufacturer’s instructions. Cells were subsequently imaged ∼24 h after transient transfection. Various stressors were used for stress granule induction in accordance with the literature. Specifically, for FUS, 0.4 M sorbitol (sigma) was dissolved directly in culture media and incubated at 37°C for 1 hour. For IL-38 cells were washed twice in media without glucose or serum and then incubated in the same media for ∼20 hours. For PKM2, cells were exposed to 500 µM sodium arsenite (Fisher) for 30–60 minutes. Laser scanning confocal microscopy was carried out using an LSM 980 (Carl Zeiss Ltd). Excitation was performed using 488 nm and 561 nm diode lasers (max. 6 mW at focal plane). The resulting fluorescence was collected using a 63× Plan-Apochromat 1.4 NA oil objective (Zeiss) and detected on a 34-channel spectral array detector in either the 510–540 nm (mEmerald) or 580–620 nm (mScarlet) range. Samples were imaged at 37°C and 5% CO_2_ in a humidified imaging chamber with a Definite Focus module (Zeiss) employed for thermal drift correction and ZEN Blue v3.9 (Zeiss) software used for acquisition. SPY650-DNA (Spirochrome) was used according to the manufacturers instructions to label Nuclei where listed in the figure legends. Here, excitation was performed with a 639 nm laser diode (max. 5 mW at focal plane) and fluorescence was collected in the 650–720 nm range.

### 6. *In vitro* imaging

To visualise lysate granules off chip, recombinant protein samples were mixed with lysates or lysis buffer to the desired concentrations as described in the figure legends. Final volumes of 25 μL of the samples were transferred to 8-well glass bottom chambered coverslips (Ibidi, 80826) and incubated at 37 °C for 3 min prior to imaging. Imaging was performed using the same set up as for live cell imaging above.

### 7. Fluorescence recovery after photobleaching (FRAP)

To observe the mobility of components within lysate granules, samples for FRAP were prepared as described above. FRAP experiments were performed on a scanning confocal LSM 780 (Zeiss) fitted with a Plan-Apochromat 100× 1.4 NA oil objective (Zeiss) at 37 °C in a humidified imaging chamber. 488 nm and 561 laser lines from a 35 mW argon multi-line laser (max. 3 mW at focal plane) were used for excitation and bleaching of the mEmerald-tagged and mScarlet-tagged components respectively. Fluorescence was detected on a 34-channel spectral array detector in the 510–540 nm (mEmerald) or 580–620 nm (mScarlet) range. Samples were stabilised for thermal drift using a Definite Focus module (Zeiss) and ZEN Black v2.3 (Zeiss) software was used for acquisition. Images were collected at 11 Hz, with 20 frames acquired prior to photobleaching, followed by an additional 330 frames to observe recovery. FRAP data were plotted using the FRAP Profiler plugin on FIJI (NIH). Briefly, the relative intensity of the photobleached region was calculated by: (i) subtracting the fluorescence background, (ii) using a non-bleached region as a control to normalize for fluorescence depletion as a result of the image acquisition, (iii) plotting the data normalised to the mean intensity of the first 20 pre-bleach frames.

### 8. Fabrication of microfluidic devices

AutoCAD software was employed for the design of the microfluidic devices, which were subsequently fabricated using conventional soft-photolithography methods. This process utilized SU8-on-Si wafer masters and polydimethylsiloxane (PDMS)-on-glass devices for production of microfluidic devices as described previously [3, 38, 39]. In short, SU-8 3050 photoresist (A-Gas Electronic Materials Limited) was poured onto a polished silicon wafer (MicroChemicals GmbH) and spun at 3000 RPM for 45 seconds using a spin coater. Afterward, SU-8 coated wafer was subjected to a soft bake process on a flat hot plate at 95°C for a duration of 15 minutes. Acetate sheet mask with the design of the device was placed on top on of SU-8 coated wafer and exposed to the UV light for 40 seconds at room temperature. Immediately following the exposure, the mask was taken off, and the SU-8 coated wafer underwent post-exposure baking on a flat hot plate at 95°C for a duration of 5 minutes. SU-8 coated wafer was immersed in a solution of propylene glycol monomethyl ether acetate (PGMEA; Sigma-Aldrich) for 10-15 minutes with intermittent agitation to remove the excessive SU-8. Lastly, the wafer was washed with isopropyl alcohol and dried by blowing with nitrogen. The master wafer for crafting microfluidic devices featuring a channel height of 50 μm was acquired. Poly(dimethylsiloxane) (PDMS; Sylgard 184 kit; Dow Corning) and crosslinking agent was mixed in a ratio of 10:1 and poured onto the master wafer kept in plastic petri dish and incubated at 60 °C for 2 h. Using a scalpel, the PDMS device was cut from the petri dish, holes of inlets and outlets were punched, and the PDMS device was cleaned in an isopropyl alcohol (IPA) bath by sonication for 15 minutes. The PDMS device was bonded onto a clean 1 mm thick glass slide (Epredia) using oxygen plasma oven (40% power for 30 s, Diener Femto Electronics). The channels of the microfluidic device were made hydrophobic by treating it with 1% v/v trichloro(1H,1H,2H,2H-perfluorooctyl)silane (Sigma) in HFE-7500 fluorinated oil (3M™ Novec™ Engineered fluid) solution for 2 minutes, and immediately dried on flat hot plate at 95 °C for 10 minutes. Finally, the channels were washed with HFE-7500 fluorinated oil and dried by blowing with nitrogen.

### 9. Generation of phase diagrams

PhaseScan, a semi-automated combinatorial droplet microfluidic platform was used to construct multidimensional 2D and 3D phase diagrams. Briefly, a 2D and 3D PhaseScan run consists of three (lysate, protein, buffer) and four (lysate, protein, buffer modulator) aqueous solutions respectively. Microfluidic flow controllers (Flow EZ^TM^, flow unit, OxyGEN, Fluigent) were used for pressure-based flow control of oil and aqueous solutions (lysate, protein, lysis buffer, modulator if present) to generate microdroplets in the microfluidic device. Each of the aqueous solutions had a unique fluorescent barcode as listed in the figure legends. A predefined flow profile was used to automatically vary the aqueous flow rates, with a total aqueous flow of 60 µL/h, maximum flow rate of 50 µL/h, minimum flow rate of 5 µL/h for a 2D PhaseScan cycle and a total aqueous flow of 80 µL/h, maximum flow rate of 65 µL/h, minimum flow rate of 5 µL/h for a 3D PhaseScan cycle to generate microdroplets with desired concentrations. HFE-7500 fluorinated oil containing 1.2 % w/v fluorosurfactant (RAN Biotechnologies) was administered into the microfluidic device at a constant flow of 10 or 20 μL/h. The microdroplets generated were imaged under continuous flow using an epifluorescence microscope (Cairn Research) equipped with 10x objective (Nikon CFI Plan Fluor 10x, NA 0.3). Image analysis was performed using a custom written Python script. Microdroplets were fitted with square bounding boxes within each image. Any erroneous detections were eliminated. The total intensity was computed from the fitted square areas, normalized to derive intensity per unit volume (based on the fitted diameter). This was then converted into concentrations by comparing the results to calibration images obtained from known barcode concentrations. Microdroplets were classified as phase separated and homogenous based on the presence and absence of condensates respectively. The phase-separated droplets were further subclassified into condensates and aggregates using an image analysis model as detailed below. The data were represented graphically as a scatter plot, showing the averaged values across conditions.

### 10. Machine Learning-assisted droplet classification

We encoded image patches containing single droplets using the activations of the last layer of a pre-trained MobileNetV2 convolutional neural network (CNN), as available in the torchvision python library. Feature embeddings were extracted for three datasets. NPM1 Model 1: dataset consisted of manually labelled droplet images, comprising 3,824 aggregates, 2,481 condensates, and 9,257 homogenous examples. NPM1 Model 2: This dataset included 3,860 aggregates, 2,524 condensates, and 9,257 homogenous examples. G3BP1 Model 3: A total of 2,748 aggregates, 2,725 condensates, and 10,924 homogenous examples were used. An XGBoost (XGB) classifier was trained on the extracted feature embeddings using the implementation provided by the ‘xgboost’ Python package (https://pypi.org/project/xgboost/). The model was trained with the following parameters: max_depth=3, eta=0.3, and objective="multi:softprob" for 100 epochs. Model performance was evaluated with ‘scikit-learn’ Python package on held-out test set comprising of 5% images sampled from respective dataset using stratified sampling. Class-weighted classification accuracy reached 97.28% for NPM1 Model 1, 97.68% for NPM1 Model 2, and 98.18% for G3BP1 Model 3. NPM1 Model 1 was trained on experiments using FUS-mEmerald wild-type and G156E lysates in the presence of NPM1 protein. NPM1 Model 2 was applied to experiments using the same lysates with NPM1 protein in the presence of either GUG RNA aptamers or UGU control sequences. G3BP1 Model 3 was trained and tested on experiments using FUS-mEmerald wild-type and R522G lysates with G3BP1 protein, also including GUG aptamer and UGU control conditions.

### 11. Microdroplet morphology-based analysis

To ensure that droplet classification relied on biomolecular condensate morphology rather than intensity-based artifacts, normalized grayscale images (0–255 pixel values) were converted into binary images (0–1) using an automated thresholding pipeline. We applied Kernel Density Estimation (KDE) to the non-zero pixel intensity distribution of each image to identify the optimal threshold separating condensate structures from background. The KDE approach estimates the probability density function of pixel intensities and uses derivative analysis to detect inflection points marking the transition from the main intensity distribution to the tail region, effectively identifying the onset of bright condensate structures. Following binary thresholding, morphological opening was applied using a 3×3 square kernel consisting of erosion followed by dilation to remove noise artifacts while closely approximating condensate morphology and area. This preprocessing pipeline eliminated intensity-dependent information while maintaining the geometric and morphological features of biomolecular condensates, ensuring that subsequent boundary detection and classification relied exclusively on shape-based characteristics rather than class-specific intensity patterns.

### 12. Dominance analysis

To extract dominance from the ExVivo PhaseScan assay, we adapted the approach described by Qian et al, 2024 [12]. The phase space was partitioned into discrete volumes defined by PolyA RNA concentration and total G3BP1 lysate concentration. For each volume, we plotted the total recombinant G3BP1 protein concentration (µM) inside the droplet against the dilute G3BP1 concentration measured in the lysate solution (mg/mL). The slope of this scatter plot, calculated using only phase-separated data points, yielded the apparent dominance D_app_, which does not represent true dominance due to unit mismatch between axes. To obtain the corrected, dimensionless dominance value, we considered a solution on the phase boundary but not yet crossed into the condensed regime. The total tagged and recombinant protein concentrations were denoted as c_R and c_T, respectively. We added a small amount of the recombinant protein Δc_R into the system, which then condensed to produce dilute and dense phase recombinant and tagged concentrations: c_dil_R, c_den_R, c_dil_T, and c_den_T. The dense phase volume was represented as v.

Mass conservation required:

(1a) (1 - v)c_dil_T + v c_den_T = c_T

(1b) (1 - v)c_dil_R + v c_den_R = c_R + Δc_R

We assumed the tagged and recombinant proteins behaved similarly, such that:

(2) c_dil_R / c_dil_T = c_den_R / c_den_T ≡ α ≫ 1

The expression for α was then:

(3) α = (c_R + Δc_R) / c_T

We focused on the dominance D, which was defined as:

(4) D = 1 - (c_dil_T + c_dil_R - c_T - c_R) / Δc_R

where (c_dil_T + c_dil_R - c_T - c_R) represented the change in dilute phase concentration upon the addition of Δc_R.

We defined the observed tagged dilute phase change as:

(5) Δc_dil_T ≡ c_dil_T - c_T

Then, the dominance was given by:

D = 1 - (c_dil_T - c_T)/Δc_R - (α c_dil_T - c_R)/Δc_R

= 1 - Δc_dil_T / Δc_R - [c_dil_T(c_R + Δc_R) - c_R c_T] / (Δc_R c_T)

= 1 - Δc_dil_T / Δc_R - (c_R / c_T) * (c_dil_T - c_T) / Δc_R - c_dil_T / c_T

= -Δc_dil_T / Δc_R * (1 + c_R / c_T) - Δc_dil_T / c_T

In the limit of a very small change Δc_dil_T → 0, the dominance was given by the negative response:

(6) D = -Δc_dil_T / Δc_R * (1 + c_R / c_T)

Which allows us to obtain the true dimensionless dominance.

### 13. RNA Aptamer sequences

The GUG aptamer sequence was 5’-UUGUAUUUUGAGCUAGUUUGGUGAC-3’, while the scrambled RNA control sequence was 5’-GGUAUUAAGCGUUAUUGUGUUGUCU-3’. Both sequences were synthesized by Biomers.net GmbH.

## Acknowledgements

We thank Matthew J Gratian and Mark Bowen of the CIMR microscopy core facility and Peter St George-Hyslop for the FUS-GFP WT and G156E proteins. We gratefully acknowledge funding from: the European Research Council (ERC) to TA and TPJK [DiProPhys 101001615] and to GK [SignAlloMod 101164135]; a Wellcome Trust Career Development Award to JNA [227745/Z/23/Z]; the Cambridge Centre for Physical Biology Pump Priming Grant to JNA and GK; the São Paulo Research Foundation [FAPESP 2023/04532-9] and [2024/05793-3] to EK and AJCF; the Swiss National Science Foundation to GP [310030_219251]; ERC ECH2020 through a Marie Skłodowska-Curie grant MicroREvolution [agreement no. 101023060] and Frances and Augustus Newman Foundation and the Finlay family grant to TS; European Molecular Biology Organization Fellowship [EMBO ALTF 349-2023] and UKRI Engineering and Physical Sciences Research Council Fellowship [EP/Z000033/1] to GC. The LSM980 in the CIMR microscopy core is supported by an MRC equipment grant [MR/Y002172/1]. Views and opinions expressed are those of the authors only and do not necessarily reflect those of the European Union or European Research Council. Neither the European Union nor the granting authority can be held responsible for them. For the purpose of open access, the authors have applied a CC BY public copyright licence to any Author Accepted Manuscript version arising.

## Data availability

All data generated or analysed during this study are included in this published article and its supplementary information files.

## Code availability

All code used in this study is available at GitHub, for PhaseScan analysis (https://github.com/rqi14/PhaseScan), Machine Learning-assisted droplet classification and morphology-based analysis (https://github.com/kjermakovs/DropletClassification),and dominance analysis (https://github.com/Rob-Scrutton/DominanceAnalysis)

## Author contributions

TA, GK, TPJK, and JNA conceived the project. TA and JNA designed the experiments. TA performed the PhaseScan assays and data analysis. FS, KJ, and TA developed and implemented the machine-learning classification pipeline. TA, JNA, EK and HC assisted with cell-based experiments and confocal imaging. SQ and EK contributed to recombinant protein purification. RS assisted with dominance analysis. PP and NP supported experimental work. AD-P and GP. provided guidance for IL-38 experiments. GC, EA, and AJCF provided resources. TS, JNA, GK, TPJK supervised the project. TPJK, GK, JNA acquired funding. TA, GK, and JNA wrote the manuscript with input from all authors. All authors reviewed and approved the final version of the manuscript.

