## Supplementary Information for "Mapping high resolution, multidimensional phase diagrams of physiological protein condensates"

[2] ariadne.ai ag, Buchrain, Switzerland

[3] Cambridge Institute for Medical Research, Department of Clinical Neurosciences, University of Cambridge, Cambridge, UK

[4] Department of Physics, Ribeirão Preto School of Philosophy, Science and Literature, University of São Paulo, Ribeirão Preto, Brazil

[5] Division of Rheumatology, Department of Medicine, Faculty of Medicine, University of Geneva, Geneva, Switzerland

[6] Department of Pathology and Immunology, Faculty of Medicine, University of Geneva, Geneva, Switzerland; Geneva Centre for Inflammation Research, Geneva, Switzerland

[7] Biophysics, Institute of Molecular Biosciences (IMB), University of Graz, NAWI Graz, Graz, Austria; Field of Excellence BioHealth, University of Graz, Graz, Austria; BioTechMed-Graz, Graz, Austria

[†] Current Address: Angiomed GmbH, Karlsruhe, Germany

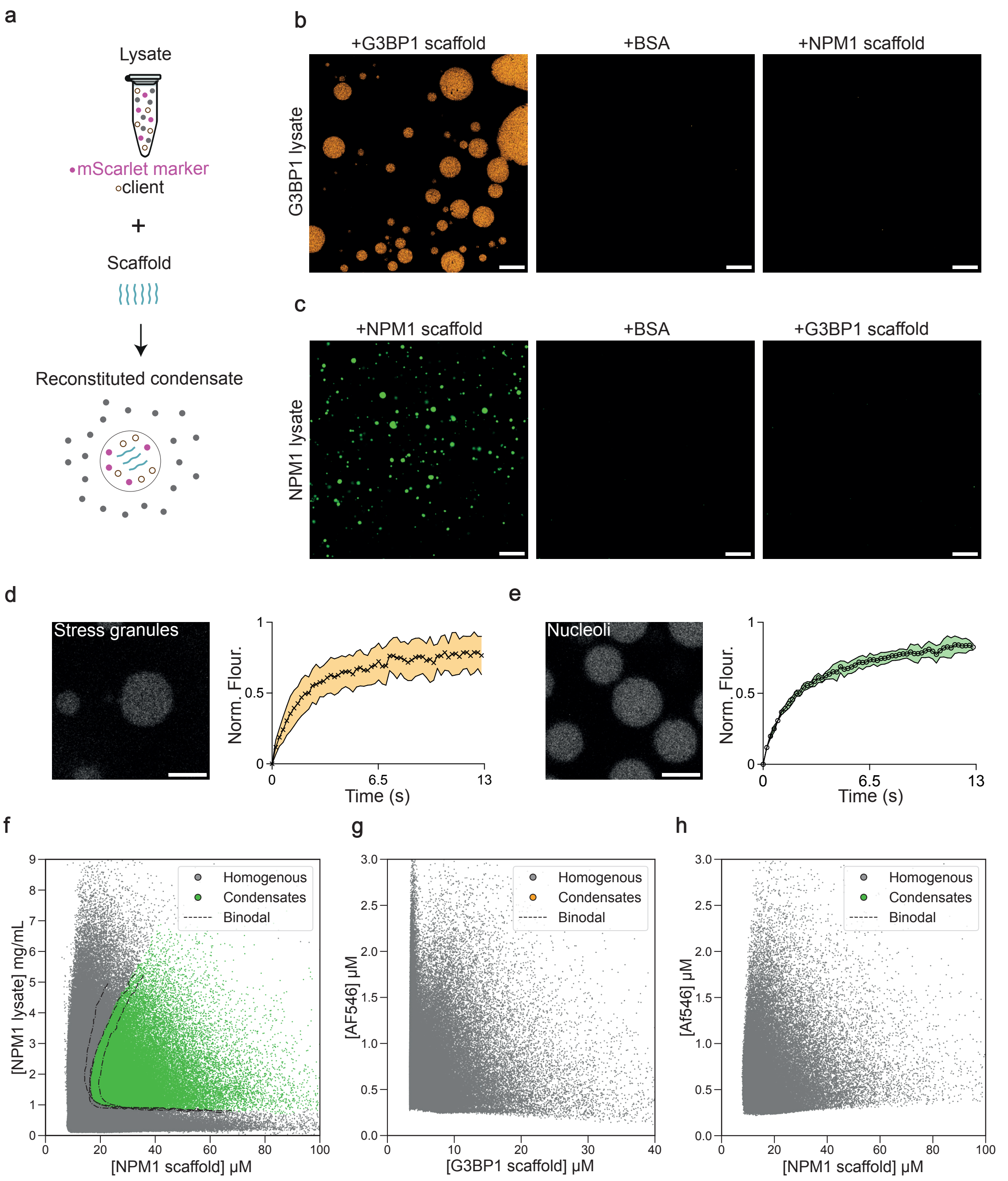


**Supplementary Figure 1: Determining the efficacy of ExVivo PhaseScan. (a)** Schematic of lysate-based reconstitution where a recombinant scaffold nucleates condensates and recruits endogenous lysate components including mScarlet-tagged markers. **(b–c)** Representative confocal images of lysate-reconstituted condensates demonstrating specificity: (b) Robust condensation with 1 mg/mL mScarlet-G3BP1 lysate and 15 μM G3BP1 scaffold (left), but no condensates with 15 μM BSA (centre) or 15 μM NPM1 scaffold (right). (c) Robust condensation with 4 mg/mL mScarlet-NPM1 lysate and 45 μM NPM1 scaffold (left), but not with 45 μM BSA (centre) or 45 µM G3BP1 scaffold (right). Scale bars, 10 µm. **(d–e)** Confocal images of reconstituted stress granules (d) and nucleoli (e) with corresponding FRAP curves, displaying fluorescence recovery. $n$ = 6 condensates, mean ± s.d. Scale bars, 20 µm. **(f)** Phase diagram of NPM1 scaffold versus NPM1-mScarlet lysate with phase boundaries from three independent experiments plotted as dashed lines, confirming reproducibility. Microdroplets are classified as homogeneous (grey) or condensate-containing (green). $n$ = 466,222 microdroplets. **(g–h)** Phase diagram of AF647 labelled G3BP1 scaffold (g) or NPM1 scaffold (h) versus AF546 barcoded lysis buffer. Microdroplets classified as homogeneous (grey) or condensate-containing (G3BP1: yellow, NPM1: green). Note the lack of condensates. $n$ = 125,528 (G3BP1); 53,693 (NPM1) microdroplets.

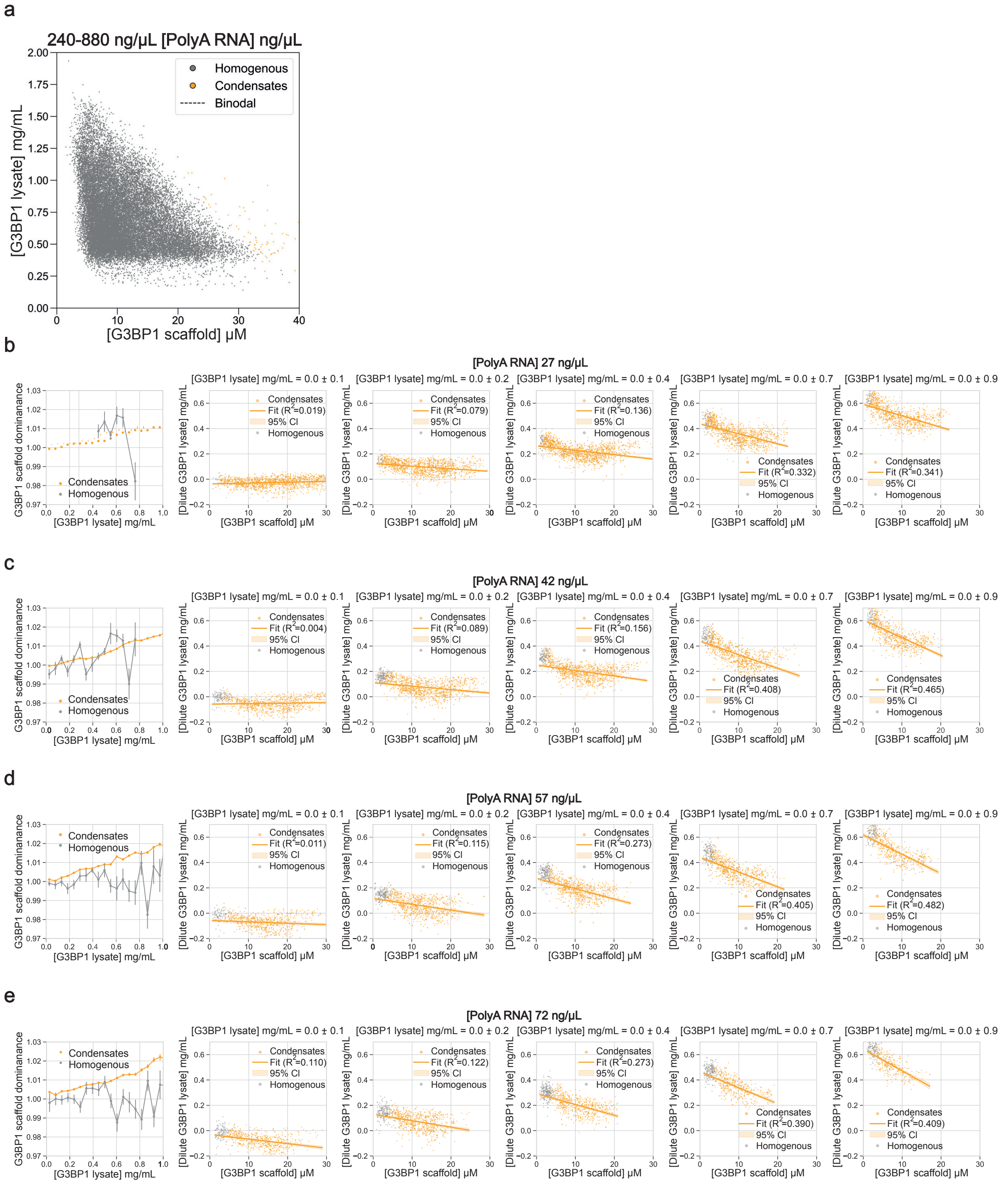


**Supplementary Figure 2: Dominance framework analysis. (a)** Phase diagram showing the effect of high Poly(A) RNA concentrations (240–880 ng/µL) on stress granule condensation. Each microdroplet is classed as homogeneous (grey) or condensate-containing (yellow). *n* = 28417 microdroplets. **(b–e)** Scatter plots showing the measured dilute phase mScarlet-G3BP1 lysate concentration (mg/mL) as a function of total recombinant G3BP1 scaffold concentration (µM) across increasing mScarlet-G3BP1 lysate concentrations (0.0–0.9 mg/mL) at four different Poly(A) RNA concentrations (27, 42, 57, and 72 ng/µL). Orange linear fits are shown with 95%

confidence intervals. The slope of each fit reflects the apparent dominance ($D_{app}$), prior to correction for unit mismatch (see Materials and Methods Section 12). Left most plots in each panel show the transition in G3BP1 scaffold apparent dominance across mScarlet-G3BP1 lysate concentration for each Poly(A) RNA level.

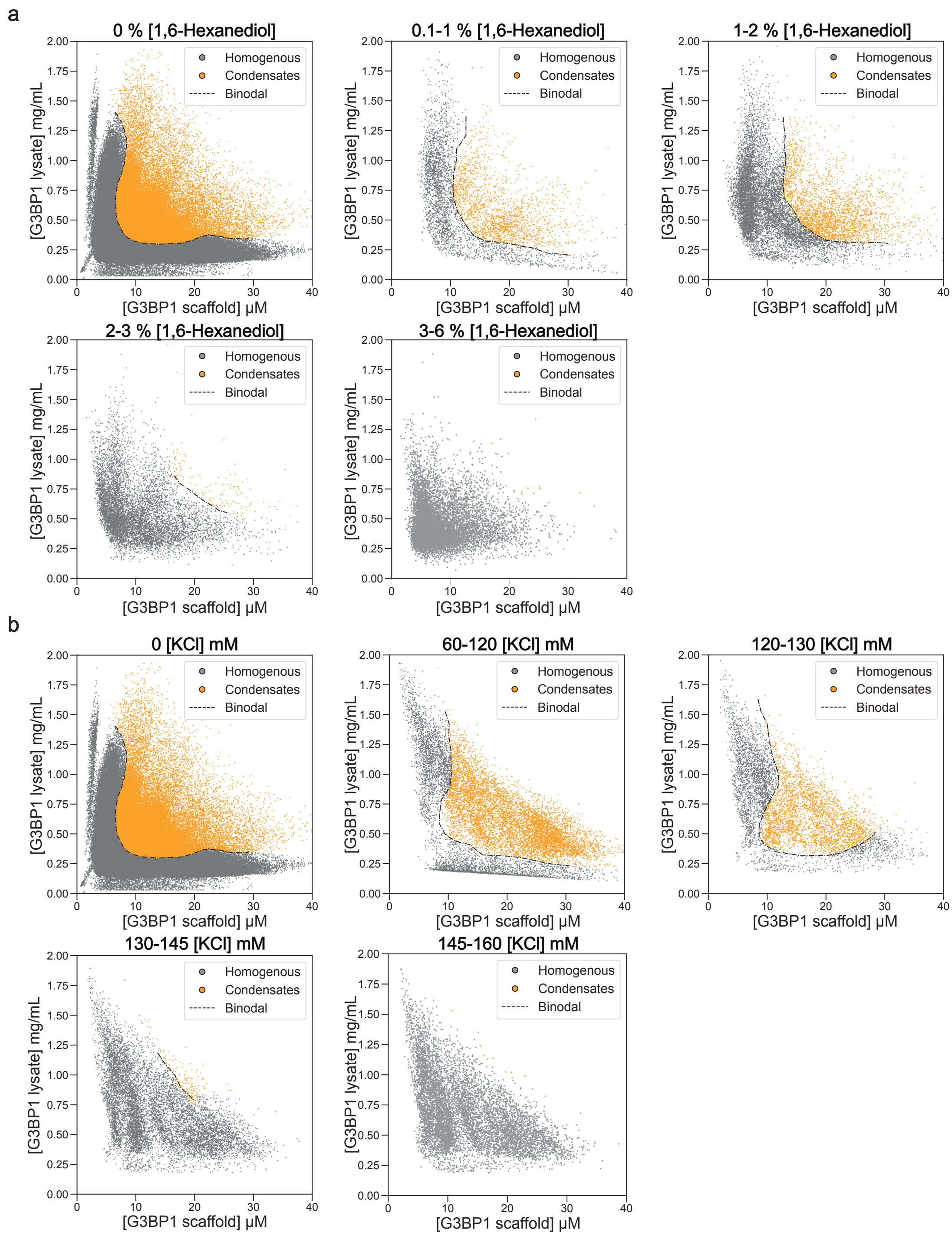


**Supplementary Figure 3: Phase diagrams showing the effect of 1,6-Hexanediol and KCl on stress granule condensation. (a)** Phase diagrams showing the effect of Increasing 1,6-Hexanediol concentrations (0%, 0.1–1%, 1–2%, 2–3%, 3–6%) on stress granule condensation. *n* = 155,584; 4,827; 11,461; 7,953; 9,361 microdroplets. **(b)** Phase diagrams showing the effect of Increasing KCl concentrations (0 mM, 60–120 mM, 120–130 mM, 130–145 mM, 145–160 mM) on stress granule condensation. *n* = 155,584; 14,314; 6,577; 9,413; 8,010 microdroplets, respectively. Microdroplets throughout are classified as homogeneous (grey) or condensate-containing (yellow), with the binodal phase boundary plotted as a dashed line.

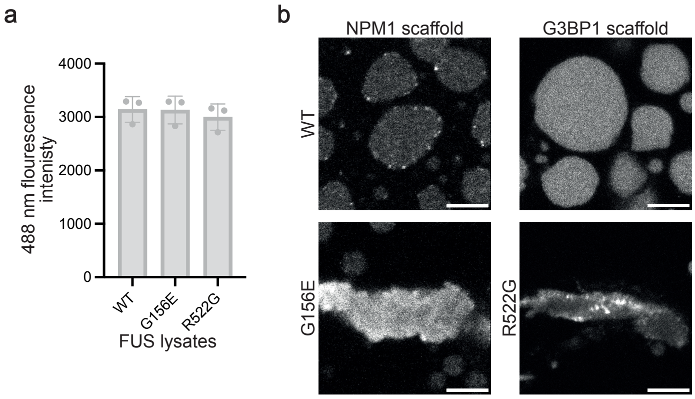


c

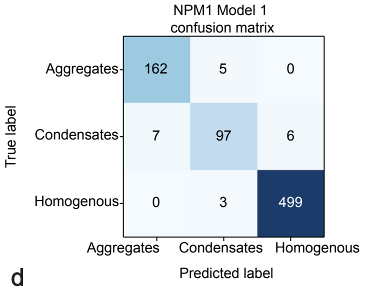


| Class | Precision | Recall | F1-Score | Support |
|---|---|---|---|---|
| Aggregate | 0.9586 | 0.9701 | 0.9643 | 167 |
| Condensate | 0.9238 | 0.8818 | 0.9023 | 110 |
| Homogenous | 0.9881 | 0.9940 | 0.9911 | 502 |
| Weighted Avg | 0.9727 | 0.9730 | 0.9728 | 779 |

d

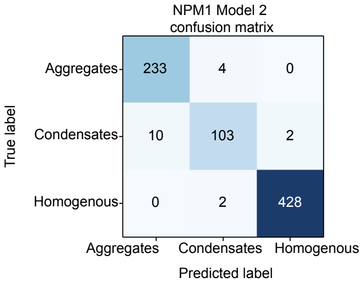


| Class | Precision | Recall | F1-Score | Support |
|---|---|---|---|---|
| Aggregate | 0.9588 | 0.9831 | 0.9708 | 237 |
| Condensate | 0.9450 | 0.8957 | 0.9196 | 115 |
| Homogenous | 0.9953 | 0.9953 | 0.9953 | 430 |
| Weighted Avg | 0.9769 | 0.9770 | 0.9768 | 782 |

e

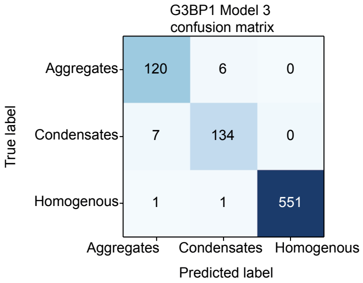


| Class | Precision | Recall | F1-Score | Support |
|---|---|---|---|---|
| Aggregates | 0.9375 | 0.9524 | 0.9449 | 126 |
| Condensates | 0.9504 | 0.9504 | 0.9504 | 141 |
| Homogenous | 1.0000 | 0.9964 | 0.9982 | 553 |
| Weighted Avg | 0.9819 | 0.9817 | 0.9818 | 820 |

**Supplementary Figure 4: Machine learning classification of FUS condensate morphology. (a)** Quantification of 488nm fluorescence intensity of prepared mEmerald-FUS lysates (WT, G156E, R522G). Mean ± s.d., $n$ = 3 independent lysate samples. One-way ANOVA with Tukey's multiple comparisons; ns ($P > 0.05$) for all groups. **(b)** Left: confocal images of reconstituted nucleoli formed using 45 μM NPM1 scaffold with 4 mg/mL mEmerald-FUS WT lysate (top) and G156E lysate (bottom). Right: confocal images of reconstituted stress granules formed using 12.5 μM G3BP1 scaffold with 5 mg/mL mEmerald-FUS WT lysate mEmerald-FUS WT lysate (top) and

R522G lysate (bottom). Scale bars-5 µm. **(c–e)** Confusion matrices and corresponding performance metrics (precision, recall, F1-score, and support) shown for three classification models used to distinguish between homogenous, condensate, and aggregate droplet morphologies. Metrics calculated on held-out test set comprising 5% of respective dataset. Weighted average scores summarize the overall performance across all classes, with each class score weighted by the number of class true instances.

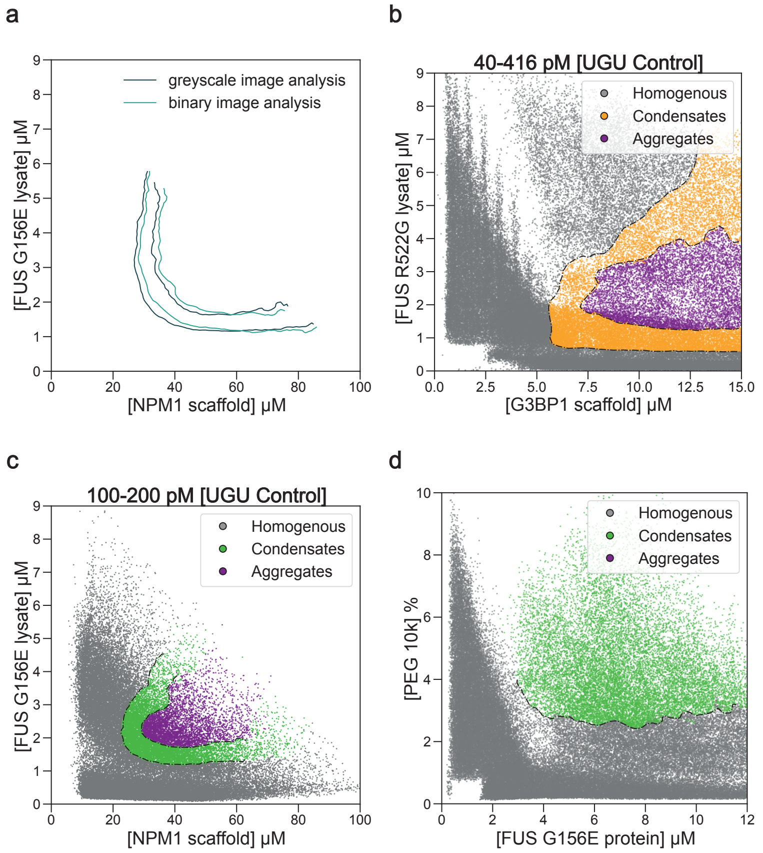


**Supplementary Figure 5: RNA aptamer-based modulation of FUS phase behaviour in stress granules. (a)** Boundary extraction from grayscale and binary image analysis (see Supplementary Text, Section 2.2). **(b–c)** Phase diagrams demonstrating the effect of a scrambled UGU control aptamer on mEmerald-FUS R522G containing stress granules (b), and mEmerald-FUS G156E containing nucleoli (c). n= 10,9195 (R522G); 81,601(G165E) microdroplets. **(d)** Phase diagram of purified G156E-GFP versus PEG 10k. *n* = 90,108; Microdroplets are classified throughout as homogeneous (grey), condensate-containing (green/yellow), or aggregated (purple), with the binodal phase boundaries plotted as a dashed line.

### Supplementary Text

#### 1.1 Dominance analysis framework

To extract the dominance from our reconstituted stress granule system, we applied the method previously outlined in Qian et al., 2024 [1] This consisted, for each individual microdroplet, calculating the total concentration and the dilute phase concentration, from the mean and 5-25% pixel intensity values respectively. Then, our 3D phase diagram was sliced into volumes of 15ng/uL RNA. In each volume, we plot each droplet's recombinant G3BP1 concentration against the dilute fluorescently tagged G3BP1-lysate concentration. The gradient of this plot for the phase separated droplets gives a response (R) related to the dominance via D = 1-R. As dominance is a dimensionless quantity, we apply a unit correction outlined in Materials and Methods Section 12. Combining the changes in dominance as a function of component concentration, with changes in the saturation concentration, we relate our observations to one of the 4 scenarios outlined in Qian et al., 2024 [1]: Target/Auxilliary Enhancement/Suppression.

#### 2.1 Machine learning-based microdroplet classifier

To enable scalable and unbiased analysis in ExVivo PhaseScan, we developed an automated machine learning pipeline to classify individual microdroplet images as homogeneous, condensate, or aggregate. Individual droplet images were first segmented, cropped, and processed through a pre-trained MobileNetV2 convolutional neural network [2] implemented via TorchVision [3]. From the final layer, we extracted 1280-dimensional feature embeddings encoding high-level spatial and textural information of the droplets. These embeddings were then used as input to an XGBoost classifier trained to distinguish three distinct morphological classes: (i) Class 1 (Homogeneous): even intensity throughout the droplet with no visible internal features; (ii) Class 2 (Condensates): presence of small in ternal puncta or dot-like patterns, indicative of liquid-like condensation; (iii) Class 3 (Aggregates): irregular internal structures consistent with misfolded or aggregated material.

#### 2.2 Microdroplet classifier validation

To demonstrate that our classification pipeline relies on condensate morphological features instead of exploiting obscure class-specific pixel intensity patterns, we compared boundary detection using per droplet normalised grayscale images (0–255 pixel values) and binarised image masks (converted to 0–1). We converted grayscale images to binary masks with a pixel value of 1 assigned to objects (condensates/aggregates) and 0 to the remaining image. Boundaries extracted from both intensity-based and binarised masks approaches are extremely similar (Supplementary Figure 5a), ensuring that our analysis relies on morphology-based features rather than class-specific information leakage. These results highlight the robustness of the pipeline in resolving subtle morphological differences and demonstrate its broad utility for high-throughput droplet phenotyping.
